# Improvement of Antibody Affinity and Using PURE Ribosome Display and Microbial Secretion System

**DOI:** 10.1101/2024.09.13.612811

**Authors:** Monami Kihara, Rio Okuda, Anri Okada, Teruyo Ojima-Kato, Hideo Nakano

## Abstract

A method has been developed for efficiently enriching and analyzing high-affinity antibody variants by combining the PURE ribosome display (ribosome display using purified cell-free protein synthesis components) with next-generation sequencing (NGS) and the *Brevibacillus choshinensis* secretion system, using the NZ-1 antibody, which targets the PA tag peptide (GVAMPGAEDDVV), as a model antibody. An artificial DNA encoding the single-chain fragment of binding (scFab) of the NZ-1 antibody was synthesized and actively expressed by cell-free protein synthesis system (CFPS). Region-specific saturation mutations were introduced into the scFab gene based on its structural information. The resulting scFab library was selected against the PA tag through two rounds of PURE ribosome display, followed by Illumina sequencing to identify potential scFab variants with higher affinity. The candidates were expressed as Fab fragments using the *B. choshinensis* secretion system. These Fab fragments were then purified from the culture supernatant using two-step column chromatography. The binding affinity of the purified Fab was evaluated using a biolayer interferometry assay, revealing a variant Fab with higher affinity than the wild-type Fab. These results demonstrate that integrating PURE ribosome display with NGS analysis and the *B. choshinensis* secretion expression system enables the rapid identification and analysis of high-affinity antibody variants.

## Introduction

Monoclonal antibodies (mAbs), which possess the ability to selectively bind to specific targets, play a crucial role in various domains such as research, biosensing, diagnostics, and pharmaceuticals^1,2^. Traditionally, mAbs have been generated using In vitro methods like the hybridoma technique or phage display^3–5^. However, advancements in technology have introduced several alternative approaches, including single B cell technology^6^, yeast surface display^7^, and In vitro display technologies like ribosome display and mRNA display/cDNA display^8^, which are increasingly employed for the selection and engineering of mAbs.

One significant advantage of the In vitro display technologies is their capacity to accommodate larger library sizes due to the elimination of transformation steps and the absence of biases associated with membrane translocation, in contrast to In vivo display methods like phage display, yeast surface display. Consequently, In vitro display technologies are getting more attention for mAb engineering. The use of an *E. coli*-based reconstituted cell-free system (PURE system), which is devoid of proteases and nucleases, further enhances the stability of peptide-ribosome-mRNA complexes, thereby facilitating the efficient selection of binding protein, named PURE ribosome display^9–11^. For instance, Fujino et al. optimized the complementarity-determining regions (CDRs) of scFabs targeting TNFaR, resulting in a remarkable 2000-fold increase in affinity (with a dissociation constant, improving from 7.28 nM to 3.45 pM)^10^.

Traditionally, enriched DNA fragments were cloned into *E. coli* to express the antibody fragments, followed by labor-intensive analysis to identify clones with the desired target properties. However, the advent of next-generation sequencing (NGS) has revolutionized this process by enabling comprehensive analysis, revealing preferred sequences in a labor-efficient manner. This technology allows for the high-throughput identification of mAb sequences with desirable characteristics, significantly reducing the time and effort required for the analysis^11,12^.

To thoroughly characterize the obtained candidate mAbs, it is essential to purify several micrograms of the candidate antibodies. Cell-free protein synthesis systems (CFPS), which allow for the straightforward synthesis of mAbs directly from PCR templates, typically yield only up to 200 ng of protein per 100 µl reaction^13^. In contrast, expressing mAbs in *E. coli* can result in the production of large quantities of protein. However, this approach often leads to the formation of insoluble proteins, requiring extensive optimization for refolding to achieve soluble mAbs with functional antigen-binding activity^14,15^.

To address these challenges, the *Brevibacillus* In vivo cloning (BIC) method was recently developed, which facilitates the secretion of large amounts of protein in a genetically engineered strain of *Brevibacillus choshinensis*^16^. This strain has been modified to enhance its protein production capabilities significantly. The BIC system has been successfully employed to produce various proteins, including scFv^16^, halophilic β-lactamase-scFv fusion proteins^17^, camelid heavy-chain antibody VHH^18^, and Fab fragments^19^, using the secretion expression system.

In this study, we utilized a sequential approach combining PURE ribosome display, NGS analysis, and the *Brevibacillus* expression system to enhance the properties of the antibody NZ-1. NZ-1 is known for its high affinity and selectivity in binding a 14-residue peptide derived from the human podoplanin PLAG domain (PA tag: GVAMPGAEDDVVV)^20,21^. Initially, based on the amino acid sequence of NZ-1 from a protein database, the encoding DNA fragment was synthesized in a single-chain Fab (scFab) form. The antigen-binding activity of the scFab synthesized using the PURE system was confirmed.

Following this, we developed a model selection system employing PURE ribosome display coupled with the SecM arrest sequence^11,22^. Region-specific saturation mutations were introduced into the scFab, guided by the structural analysis of the NZ-1 antibody-PA tag peptide complex. To identify scFab variant s with higher affinity, we performed two rounds of selection using the ribosome display system, followed by NGS analysis. Several promising variant candidates were expressed using the *Brevibacillus* system, purified, and their affinities were analyzed. The results indicated that one Fab variant exhibited a higher affinity than the wild-type scFab.

We will discuss the feasibility of this integrated process for enhancing antibody affinity and achieving cost-effective microbial production for industrial applications.

## Methods

### 1. Plasmid construction

For producing PA tag antigen fused with maltose-binding protein (MBP), we first constructed the MBP-PA expression plasmid. PCR was performed by PrimeSTAR Max (Takara Bio Inc., Japan) using the plasmid pMal-c2-MalE-XlnR1-183^23^, which contains MBP, as the template with the primers pmalc2-patag-F and pmalc2-patag-R, both of which collectively encode the PA tag (GVAMPGAEDDVVV), under the following conditions: initial denaturation at 94°C for 15 seconds, followed by 25 cycles of denaturation at 94°C for 15 seconds, and annealing and extension at 68°C for 7 minutes and 20 seconds. The PCR product was self-ligatated by SLiCE.^24^ *E. coli* DH5α was transformed by the SLiCE reaction mixture and plated on LB agar containing 100 µg/ml ampicillin. A single colony was selected and used as template for colony PCR with primers (F1 and R1). The amplified DNA was sequenced to confirm the whole of the MBA-PA protein. The same colony was also inoculated into LB medium with 100 µg/ml ampicillin and cultured overnight at 37°C and 110 rpm. The pMalc2-MBP-PA plasmid was extracted using the FastGene Plasmid Mini Kit (Nippon Genetics Co. Ltd, Japan).

The plasmid pRSET-anti-PAscFab, which includes the His tag and SecM arrest sequence (A148-T170), was constructed as follows (Figure 1A). The amino acid sequences of the heavy and light chain variable regions of the antibody NZ-1, which has high affinity for the PA tag, were obtained from the database (PDB : 6ICF). The nucleotide sequences were designed based on *E. coli* codon usage. The designed DNA sequences were synthesized as gene block fragments (Integrated Gene Technologies Inc, USA). The mutant plasmid pRSET-anti-PAscFab(118-120Ala) was created by replacing the amino acids Thr118, Ser119, and Arg120 in the heavy chain, which are important for binding the antigen PA tag, with Ala.

**Figure 1.**
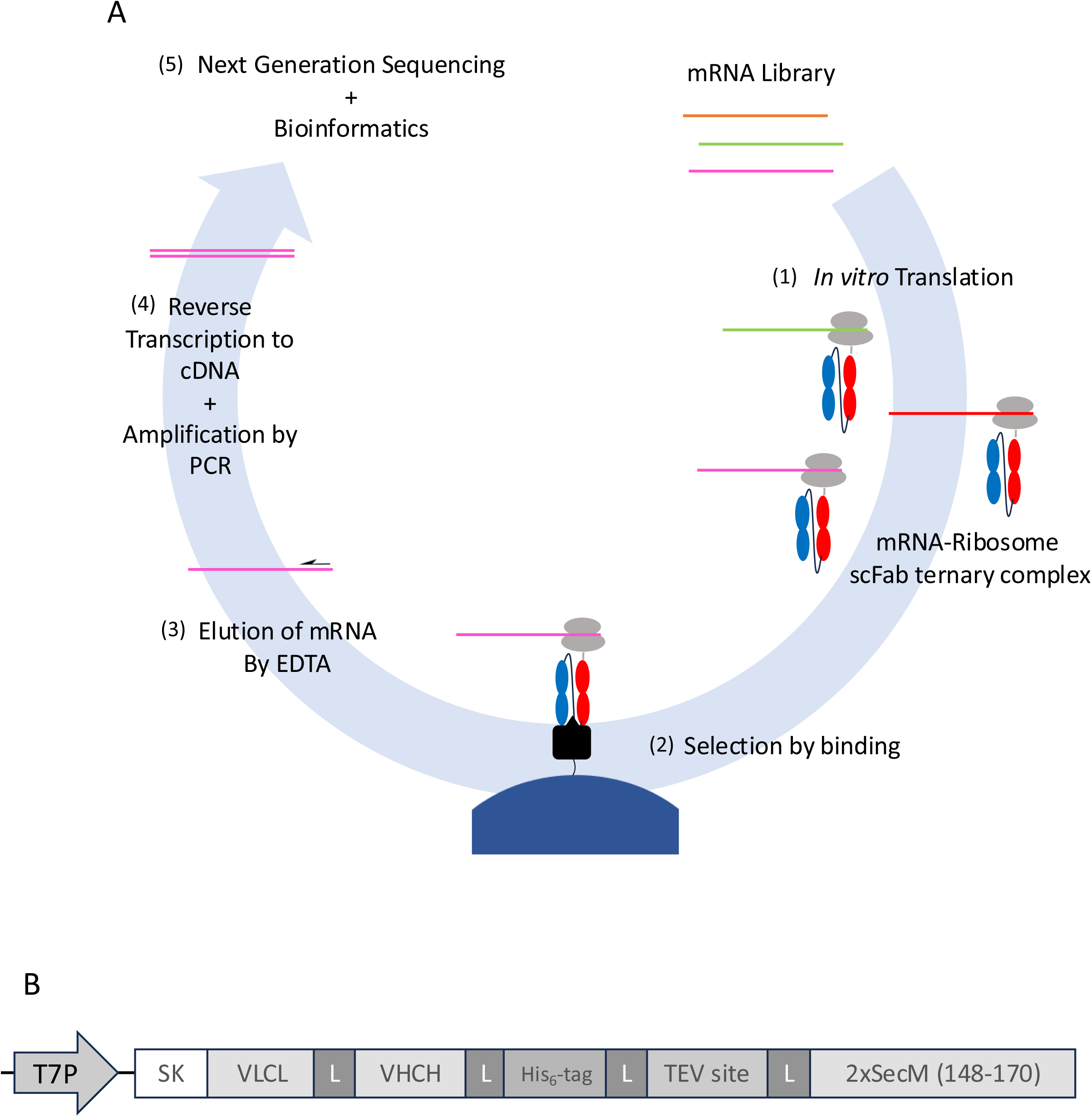
Overview of the PURE ribosome display platform for anti-PA tag scFab selection A : Schematic protocol of the PURE ribosome display platform. (1) Formation of protein–ribosome-mRNA ternary complexes through translation of the mRNA library using the PURE system. (2) Selection of ternary complexes that bind to the antigen. (3) Separation of mRNA from the selected ternary complexes. (4) Preparation of a DNA pool for NGS analysis using RT-PCR. (5) Analysis of enriched scFab sequences using NGS data. B : The DNA template for ribosome display.

**Figure 2.**
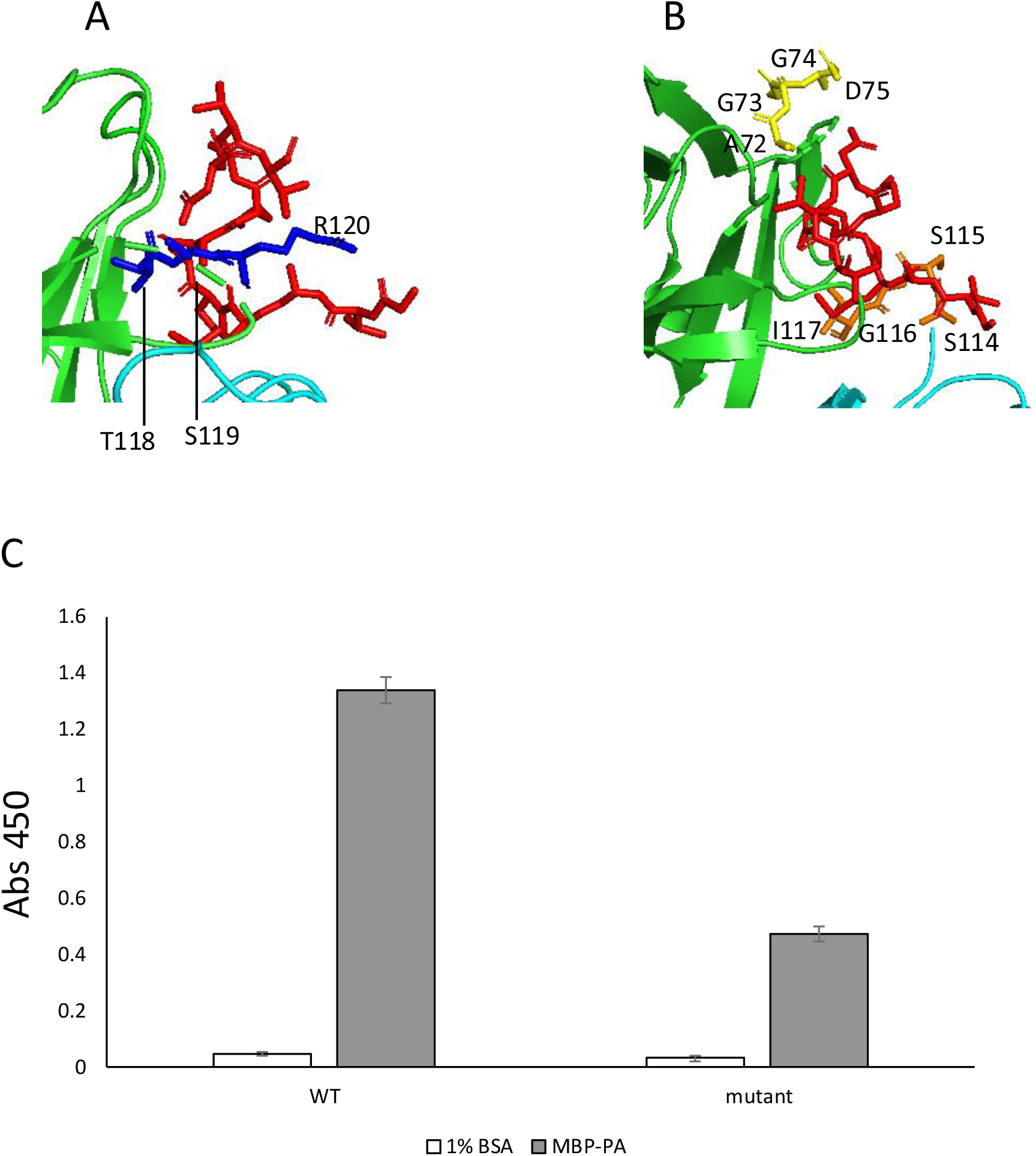
Structural and mutational analysis of anti-PA tag fab and NZ-1 A: Locations of alanine mutations introduced to create a low-binding mutant scFab (anti-PAscFab(118-120Ala)) from anti-PA tag Fab (NZ-1). 3D structure of NZ-1 (PDB: 6ICF) is shown as Green: Heavy chain (Hc), Cyan: Light chain (Lc), Red: PA tag, Blue: amino acid residues (Thr118, Ser119, Arg120) substituted to Ala. B: Locations of region-specific mutations introduced Lc library regions S114, S115, G116, I117 are shown in orange and Hc library regions A72, G73, G74, D73 are shown in yellow. C: ELISA results of the anti-PAscFab and anti-PAscFab(118-120Ala). The anti-PAscFab DNA and the anti-PAscFab(118-120Ala) DNA were expressed by CFPS with *PUREflex2*.1 (GeneFrontier, Japan). Wells pre-coated with 1% BSA or MBP-PA were incubated with 100-fold diluted CFPS products. X-axis: ELISA signal (Abs 450); error bars: standard deviation of the mean (n=7).

The plasmid pRSET-anti-PAscFab was used as template for PCR amplification by KOD Fx Neo with the primers LCVector_F_60 and LCVector_R_60 under the following conditions: initial denaturation at 94°C for 1 minute, followed by 25 cycles of denaturation at 94°C for 10 seconds, annealing for 15 seconds at 61°C, and extension for 4 minutes at 68°C. The amplified products were purified, self-ligated and transformed *E. coli* DH5α, resulting in the construction of a plasmid pRSET-anti-PA-LCSSGI, which lacks Ser114-Ile117.The procedure for constructing pRSET-anti-PA-LCSSGI was similar to that used to construct pRSET-anti-PA-HCSA. The template pRSET-anti-PAscFab was amplified by KOD Fx Neo with primers HCVector_F_58 and HCVector_F_58. The amplified products were purified. Ligation reactions were performed using the SLiCE method, and the plasmid pRSET-anti-PA-HCSA, which lacks Ala72-Asp75, was extracted.

### 2. Expression and Purification of MBP-PA

*E. coli* BL21 (DE3) cells were transformed with the pMalc2-MBP-PA plasmid and cultured on LB agar containing 100 µg/ml ampicillin. A single colony was selected and inoculated into LB medium supplemented with 100 µg/ml ampicillin, followed by overnight incubation at 37°C with shaking at 110 rpm. This culture was then transferred to 100 ml of fresh LB medium and grown under the same conditions. Upon reaching an OD600 of 0.5, IPTG was added to a final concentration of 1 mM, and induction was carried out at 30°C and 110 rpm for 18 hours.

After induction, the culture was centrifuged at 9000 rpm for 10 minutes at 4°C, and the resulting cell pellet was resuspended in 25 ml of Sonication Buffer (200 mM NaCl, 20 mM Tris-HCl pH 7.5). The cells were then sonicated on ice for 40 minutes. The lysate was clarified by centrifugation at 9000 rpm for 10 minutes at 4°C, and the soluble fraction was collected.

The soluble fraction was diluted with 150 ml of Binding Buffer (200 mM NaCl, 20 mM Tris-HCl, 1 mM EDTA, 10 mM β-mercaptoethanol, pH 7.5) and subjected to purification using 1 mL of Amylose Resin (New England Biolabs, Japan). The column was equilibrated with 5 column volumes (CV) of Binding Buffer before loading the diluted sample. After washing with 5 CV of Binding Buffer, the bound protein was eluted with 3 CV of Elution Buffer (200 mM NaCl, 20 mM Tris-HCl, 1 mM EDTA, 10 mM β-mercaptoethanol, 10 mM maltose). The eluted protein fraction was dialyzed overnight at 4°C against Dialysis Buffer (10 mM Na_2_HPO_4_·12H_2_O, 1.8 mM KH2PO4, 137 mM NaCl, 2.7 mM KCl, 1 mM β-mercaptoethanol, 30% glycerol). The purified MBP-PA was then collected and stored at - 20°C.

### 3. Preparation of MBP-PA Conjugated Magnetic Beads

The purified MBP-PA was biotinylated using the Biotin Labeling Kit-NH2 (Dojindo Laboratories Co., Ltd., Japan) and stored at 4°C. A hundred µl Dynabeads M-280 Streptavidin (Thermo fisher scientific, USA) were washed with 1 ml of PBS, followed by the addition of 100 µl of biotinylated MBP-PA. The mixture was incubated with rotation at room temperature for 1 hour. The beads were then washed with 1 ml of PBS to yield MBP-PA conjugated M-280 Streptavidin Magnetic Beads (MBP-PA magnetic beads).

### 4. ELISA with scFab synthesized by cell-free protein synthesis

To compare anti-PAscFab and anti-PAscFab(118-120Ala), an ELISA was conducted using scFab proteins translated from scFab DNA pools via CFPS. The DNA fragments used in the PURE ribosome display were reamplified with F1 and NoSecM-Rv primers to remove the SecM sequence. The anti-PAscFab DNA and anti-PAscFab(118-120Ala) DNA were used as templates for CFPS using PURE*frex* 2.1 (GeneFrontier, Japan). 96-well plate (Thermo Fisher Scientific) coated with 1% BSA or 10 µg/ml of MBP-PA was incubated at 37℃ for 2 hours. Then the plates were washed twice with phosphate-buffered saline (PBS) and blocked with 1% skim milk (Nacalai Tesque Inc., Kyoto, Japan) in PBS at room temperature for 1 hour. The resulting CFPS reactions were diluted 1:100 with Can Get Signal Solution 1 (TOYOBO, Japan) and then added to the wells and incubated at room temperature for 1 hour. The plate was washed twice with PBST (PBS with 0.05% Tween-20). Anti-His-tag mAb-HRP-DirecT (MBL Co. Ltd., Japan) diluted 1:5000 with Can Get Signal Solution 2 (TOYOBO, Japan) was added and incubated at room temperature for 1 hour. The plate was washed three times with PBST and 1-step Ultra TMB-ELISA Substrate Solution (Thermo fisher scientific, USA) was added, incubating at room temperature for 10 minutes. The reaction was stopped with 2 mol/l sulfuric acid and the absorbance (Abs) at 450 nm was measured using a plate reader (Infinite M200 PRO, TECAN Japan, Japan).

### 5. PURE Ribosome Display with Model Library

pRSET-anti-PAscFab and pRSET-anti-PAscFab(118-120Ala) plasmids were used as templates for PCR amplification with F1 and RT-R primers. The amplified products were purified using the FastGene™ Gel/PCR Extraction Kit (Nippon Genetics Co. Ltd, Japan). The purified DNA templates were transcribed using the RiboMAX Large Scale RNA Production System-T7 (Promega, USA) at 37°C for 2 hours, treated with DNase I (Takara Bio Inc, Japan) at 37°C for 30 minutes to remove the template DNA followed by mRNA purification using the RNeasy® MinElute® Cleanup Kit (Thermo fisher scientific, USA). The purified mRNA was quantified using Nanodrop One (Thermo fisher scientific, USA). The obtained two kinds of mRNA were mixed at a ratio of anti-PAscFab : anti-PAscFab(118-120Ala) = 1:100 (1 ng : 100 ng).

The mixed mRNA was added to the PURE*frex* 1.0 cell-free protein synthesis system (GeneFrontier, Japan) at a 20 µl reaction scale and incubated at 30°C for 30 minutes to form the scFab-ribosome-mRNA ternary complex. The reaction mixture was diluted 10-fold with ice-cold WBTBR (50 mM Tris-HCl pH 7.6, 90 mM NaCl, 50 mM Mg(OAc)_2_, 0.5% Tween20, 5 mg/ml BSA (Sigma), 1.25 mg/ml yeast RNA (Sigma), 0.04 U/µl Recombinant RNase Inhibitor (Takara Bio Inc., Japan). The reaction mixture was mixed with 100 µl of MBP-PA magnetic beads and rotated at 37°C for 1 hour. The MBP-PA magnetic beads were washed three times with WBT (50 mM Tris-HCl pH 7.6, 150 mM NaCl, 50 mM Mg(OAc)_2_, 0.05% Tween-20). Elution Buffer (50 mM Tris-HCl pH 7.6, 150 mM NaCl, 50 mM EDTA) was added to the MBP-PA magnetic beads and the supernatant was collected. Reverse transcription was performed on the collected supernatant using SuperScript IV reverse transcriptase (Thermo fisher scientific, USA) and RT-Rv primer. The reverse-transcribed cDNA was amplified using KOD One PCR Master Mix (TOYOBO, Japan) and PCR1-Fw/Rv primers under the following conditions: initial denaturation at 94°C for 10 seconds, followed by 30 cycles of denaturation at 94°C for 15 seconds, and annealing and extension for 4 minutes at 68°C. The anti-PAscFab and anti-PAscFab(118-120Ala) were distinguished by utilizing a *Sac*II restriction site present only in the anti-PAscFab(118-120Ala) DNA. The PCR products were treated with *Sac*II (Takara Bio Inc, Japan) and analyzed by 1% agarose gel electrophoresis.

### 6. Construction of Anti-PA Tag Antibody Random Library

Based on the crystal structure of the complex with the antigen PA12 tag (PDB ID: 6ICF), four amino acid residues (Ser114, Ser115, Gly116, and Ile117) in the light chain CDR, which are close to the PA tag peptide, were identified and replaced with NNK codons to construct the light chain (Lc) library. Similarly, four amino acid residues (Ala72, Gly73, Gly74, Asp75) in the heavy chain (Hc), located near the PA tag, were replaced with NNK codons to construct the Hc library.

Oligo DNA primers LCMutant_F and HCMutant_F, containing the four NNK codons, along with dNTPs and 10x Klenow Fragment (Takara Bio Inc, Japan), were mixed and heated in boiling water for 5 minutes. The mixture was then allowed to cool to room temperature, followed by incubation at 37°C for 1 hour with 2 µl of Klenow Fragment (Takara Bio Inc, Japan) and inactivation at 65°C for 5 minutes, resulting in the synthesis of random four NNK DNA libraries covering all 20 amino acids.

The plasmid pRSET-anti-PA-LCSSGI was used as the template to amplify the vector for Lc library, using KOD Fx Neo with primers LCVeclib_F and LCVeclib_R under the following conditions: an initial denaturation at 94°C for 1 minute, followed by 25 cycles of 94°C for 10 seconds, annealing at 61°C for 15 seconds, and extension at 68°C for 4 minutes, followed by DNA purification. The vector for Hc library was similarly created using the template plasmid pRSET-anti-PA-HCSA, amplified by KOD Fx Neo with the primers HCVeclib_F and HCVeclib_R under the following conditions: initial denaturation at 94°C for 1 minute, followed by 25 cycles of 94°C for 10 seconds, annealing at 60°C for 15 seconds, and extension at 68°C for 2 minutes, followed by DNA purification.

The Lc library and the Hc library DNA fragments were ligated to the vectors for Lc library and Hc library, respectively, using the NEBuilder HiFi DNA Assembly (New England Biolabs, Japan) at 50°C for 15 minutes. The reaction mixtures were used as templates for PCR amplification with F1 and RT-R primers, followed by DNA purification. The purified DNA templates were transcribed using the RiboMAX Large Scale RNA Production System-T7 (Promega, USA) at 37°C for 2 hours, treated with DNase I (Takara Bio Inc, Japan) at 37°C for 30 minutes to remove the template DNA followed by mRNA purification using the RNeasy® MinElute® Cleanup Kit (Thermo fisher scientific, USA). The purified mRNA was quantified using Nanodrop One (Thermo fisher scientific, USA).

### 7. PURE Ribosome Display with Random Library

Two kinds of mRNA synthesized from Lc and Hc library were used as template for 20 µl cell-free protein synthesis (CFPS) using PURE*frex* 1.0, resulting in the formation of the scFab-ribosome-mRNA ternary complex. The resulting CFPS reactions were used for PURE ribosome display as described above.

For further selection, the purified DNA fragments were ligated by Gibson assembly (New England Biolabs, Japan) with DNA fragments amplified by PCR from the vector plasmid for the library with the primers Gibvec_F and GibVec_R under standard conditions. The assembled DNAs were then amplified using F1 and RT-R primers to generate DNA templates for additional rounds of ribosome display selection. The second round of PURE ribosomal display selection was performed under the same conditions of the first-round selection.

To evaluate the scFab selection against the PA tag, an ELISA was conducted using scFab proteins translated from scFab DNA pools via CFPS. The assembled DNA used in the PCR to amplify DNA templates for the second round of PURE ribosome display were reamplified with F1 and NoSecM-Rv primers to remove the SecM sequence. The selected mutant DNA pools were used as template for CFPS using PURE*frex* 2.1 (GeneFrontier, Japan). The resulting CFPS reactions were used for ELISA as described above.

### 8. Preparation of NGS Samples

The DNA pool obtained from the ribosome display was amplified using F1 primers and barcode primers (LcBf/LcAf/Lc-Rv/HcBf/HcAf/Hc-Rv). The amplified products were then purified, and equal amounts of the purified products were combined to prepare the NGS samples. The quality of the prepared samples was evaluated using the High Sensitivity D5000 ScreenTape System (Agilent). Once the quality was confirmed, the samples were pooled for next-generation sequencing, which was performed at Nagoya University’s Genetic Research Center using the Illumina NextSeq550 platform.

### 9. Analysis of NGS Sequence Data

The random regions of the library sequences were extracted from the raw Illumina sequencing data using barcode sequences and SeqKit^25^. The extracted nucleotide sequences were then translated into amino acid sequences. Sequences containing stop codons were filtered out using the R programming language, and the remaining sequences were counted for each of the four amino acids.

The enrichment rate of each sequence of the four amino acids was calculated by determining the frequency of each sequence in the total number of reads before and after selection. The enrichment factor (EF) was calculated as the ratio of the frequency after selection to the frequency before selection.

### 10. Energy Calculation

The energy minimization of the complex of antibodies was carried out using GROMACS based on the crystal structure of NZ-1 (PDB: 6ICF). The single scFab variants were predicted by AlphaFold2. To solve the system, TIP3P water was used to fill a cubic box, and the protein was placed at the center with 10 angstrom minimum distance to the box edge. Na^+^ and Cl^−^ ions were introduced to neutralize the protein charge and simulate a salt solution with a concentration of 0.15 M. Systems were minimized for 5000 steps with the steepest descent algorithm. After minimization the potential energy change of the system was calculated using the GROMACS analysis package.

### 11. Large-Scale Production and Purification of Fab Using *Brevibacillus choshinensis*

*Brevibacillus choshinensis* expression plasmids for the mutants obtained from the NGS sequence data analysis were constructed as a shuttle vector between *E.coli* and *B. choshinensis.* The antibody sequences were cloned into the pNCMO2 (Takara Bio Inc, Japan) vector to create plasmids pNCMO2_Anti-PAtag_Fab_His, pNCMO2_Anti-PAtag_A72D_Fab_His, pNCMO2_Anti-PAtag_G74C_Fab_His, pNCMO2_Anti-PAtag_G74S_Fab_His, and pNCMO2_Anti-PAtag_D75N_Fab_His. The signal peptide sequences for VL-CL and VH-CH were derived from pBIC3 and pBIC4 (Takara Bio Inc, Japan), respectively.

*Brevibacillus choshinensis* competent cells (Takara Bio Inc, Japan) were transformed with the plasmids and plated on MTNm agar medium (10 g/l phytone peptone(Thermo fisher scientific, USA), 10 g/l glucose, 5.75 g/l 35% Ehrlich bonito extract (Kyokuto Pharmaceutical, Japan), 2 g/l bacto yeast extract (Thermo fisher scientific, USA), 10 mg/l FeSO4·7H_2_O, 8.9 mg/l MnCl_2_·4H_2_O, 1 mg/l of ZnSO_4_·7H_2_O, 50 µg/ml neomycin, 15 g/l Agar). A single colony was inoculated into TM medium containing 50 µg/ml neomycin and cultured overnight at 37°C with shaking at 110 rpm. An aliquot (10 µl) of the culture was transferred to a deep-well plate containing 990 µl of TM medium. The plate was sealed with a gas-permeable membrane and incubated at 30°C with shaking at 1000 rpm for 48 hours. The culture was then centrifuged (20,000 g, 4°C, 10 minutes) and the supernatant was collected and filtered through a 0.8 µm filter.

For purification of proteins, the supernatant was dialyzed overnight at 4°C against dialysis buffer (100 mM Tris-HCl pH 7.0, 150 mM NaCl). The dialyzed supernatant was then diluted with 2 volumes of binding buffer (100 mM Tris-HCl pH 7.0, 500 mM NaCl, 10 mM Imidazole) and incubated with 2 ml of Ni-NTA Agarose (Fujifilm Wako Pure Chemical Corporation, Japan) on ice for 1 hour. After centrifugation at 500 g for 5 minutes at 4°C, the supernatant was discarded, and the resin was transferred to a disposable plastic column (Thermo fisher scientific, USA). The column was washed three times with 6 ml of wash buffer (100 mM Tris-HCl pH 7.0, 500 mM NaCl, 20 mM Imidazole). The bound protein was eluted with 6 ml of elution buffer (100 mM Tris-HCl pH 7.0, 500 mM NaCl, 300 mM Imidazole). The eluted fractions were dialyzed overnight at 4°C against dialysis buffer (100 mM Tris-HCl pH 7.0, 500 mM NaCl) and the resulting products were used for ELISA as described above.

Further purification was achieved using size-exclusion chromatography (SEC) on a HiPrep 16/60 Sephacryl S-200 HR (Cytiva, Japan) equilibrated with SEC buffer (50 mM Tris-HCl pH 7.5, 200 mM NaCl). The Fab solution was applied and eluted with 1 column volumes (CV) of SEC buffer. The purified Fab was collected and stored at −80°C. Subsequently, 7 µl of the purified Fabs were mixed either with 1 µl of1M DTT or 1 µl of water, and 2 µl 5×sample buffer (10 w/v% SDS, 50 v/v% glycerol, 30 mM Tris–HCl pH 6.8, 0.05 w/v% bromophenol blue) and boiled for 10 minutes. The samples were subjected to SDS-PAGE (12.5% acrylamide gel) and CBB staining.

### 11. Measurement of binding kinetic of Fab

The binding properties of the Fab to PA tags were analyzed using biolayer interferometry (BLI) on the BLItz system (Sartorius Japan, Japan). The APS biosensor was hydrated in PBS for 10 minutes prior to use. Each measurement commenced with a 30-second baseline measurement in PBS. Subsequently, the APS biosensor was immersed in a 50 µg/ml MBP-PA solution for 120 seconds to immobilize the antigen on the biosensor surface. This was followed by a 30-second baseline measurement in PBS. The APS biosensor was then immersed in the Fab solution for 300 seconds to measure the association rate. Finally, the sensor was transferred to PBS for 300 seconds to monitor the dissociation rate. The binding curves were generated using the BLItz Pro software, and association rate, dissociation rate, and dissociation constant were subsequently calculated.

## Results

### 1. Development of PURE ribosome display using model library

The anti-PA tag single-chain Fab (anti-PAscFab) gene, synthesized based on the amino acid sequence of the anti-PA tag antibody NZ-1, and its mutant scFab (anti-PAscFab(118-120Ala)), as described in the Materials and Methods, were used as templates for CFPS. Both scFabs were successfully expressed, as confirmed by western blotting (data not shown). ELISA analysis of the scFabs demonstrated that anti-PAscFab(118-120Ala) exhibited significantly lower binding activity compared to anti-PAscFab as shown in Figure 3. Consequently, we attempted to establish ribosome display conditions using anti-PAscFab and anti-PAscFab(118-120Ala).

**Figure 3.**
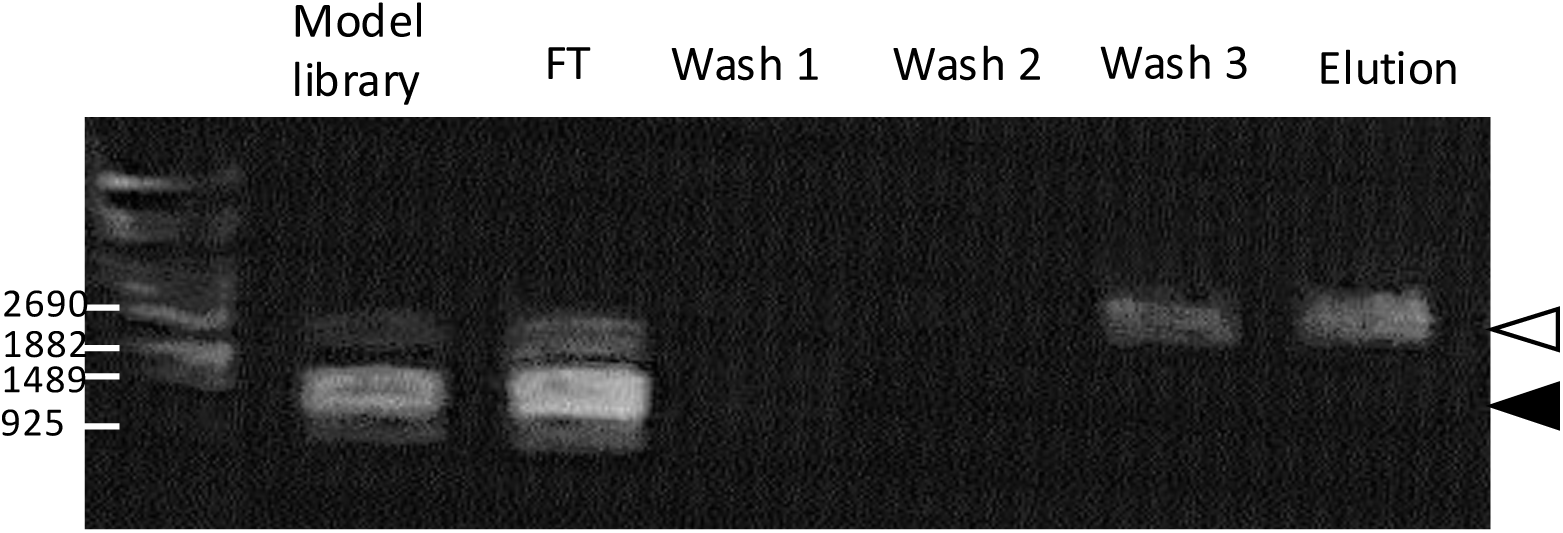
Enriched DNA analysis after one round of PURE ribosome display selection for the model library. Agarose Gel Electrophoresis results of *Sac*II-treated DNA samples from the input model library and the DNA pools from each steps of PURE ribosome display as in the section 5 of Materials and Methods. Model library: The DNA library mixed in a 1:100 ratio of anti-PAscFab DNA to anti-PAscFab(118-120Ala) DNA. FT: Flow-through; Wash1-3: First to third washes; Elution: DNA obtained from selection. All DNA samples were treated with *Sac*II and then subjected to electrophoresis. The white triangle indicates the size of anti-PAscFab, the black triangle indicates the size of anti-PAscFab(118-120Ala).

To form a stable protein-ribosome-mRNA ternary complex, the SecM peptide sequence, which induces ribosome stalling and stabilizes the complex, was fused to the C-terminus of each scFab. A model library was prepared with a 1:100 molar ratio of mRNA from anti-PAscFab : anti-PAscFab(118-120Ala), and the translation was carried out using CFPS. The resulting ternary complexes were captured by MBP-PA magnetic beads. After one round of selection, approximately 90% of the DNA from the model library corresponded to anti-PAscFab (Figure 3), indicating that ribosome display is effective under these conditions.

### 2. Enrichment of high affinity scFab by PURE ribosome display from region-specific random library

To obtain variants anti-PA tag scFabs with enhanced binding abilities using ribosome display, we constructed two libraries: the Lc library and the Hc library as described in Materials and Methods. Ribosome display was performed against the PA tag using these libraries. scFabs were synthesized from the DNA pools before and after selection using transcription and translation with the PURE system, and their binding abilities to the antigen were analyzed via ELISA. After selection, the ELISA signal for MBP-PA increased by 1.1-fold in the Lc library and 2.5-fold in the Hc library, compared to pre-selection (Figure 4).

**Figure 4.**
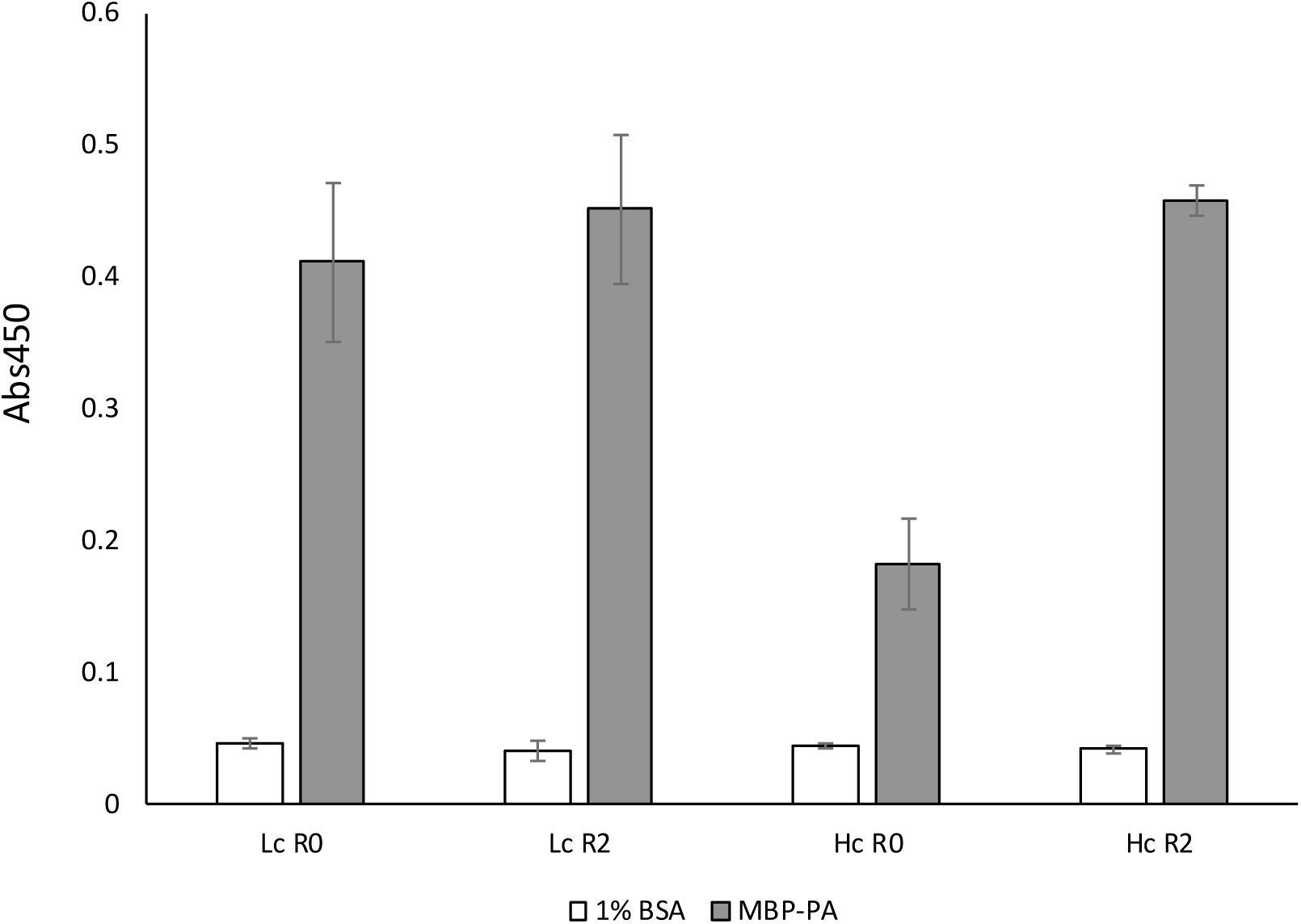
ELISA analysis to compare with pre- and post-selection DNA pools. The selected DNA pools (R2) and the initial library (R0) were expressed by CFPS with *PUREflex 2*.1. Wells pre-coated with 1% BSA or MBP-PA were incubated with 100-fold diluted CFPS products. CFPS was performed using pre-selection DNA pools (Lc R0 and Hc R0) and DNA pools post-selection (Lc R2 and Hc R2) as templates. X-axis: ELISA signal (Abs 450); error bars: standard deviation of the mean (n=6).

In the selection using the Hc library, the wild-type sequence, AGGD, exhibited a high enrichment rate, with an EF of 93.32. Additionally, other sequences such as AGCD, AGSD, DGGD, and AGGN showed similarly high EF (Table 1). On the other hand, in the selection of the Lc library, the wild-type sequence, SSGI and SSCI had significantly high EF.

**Table 1.**
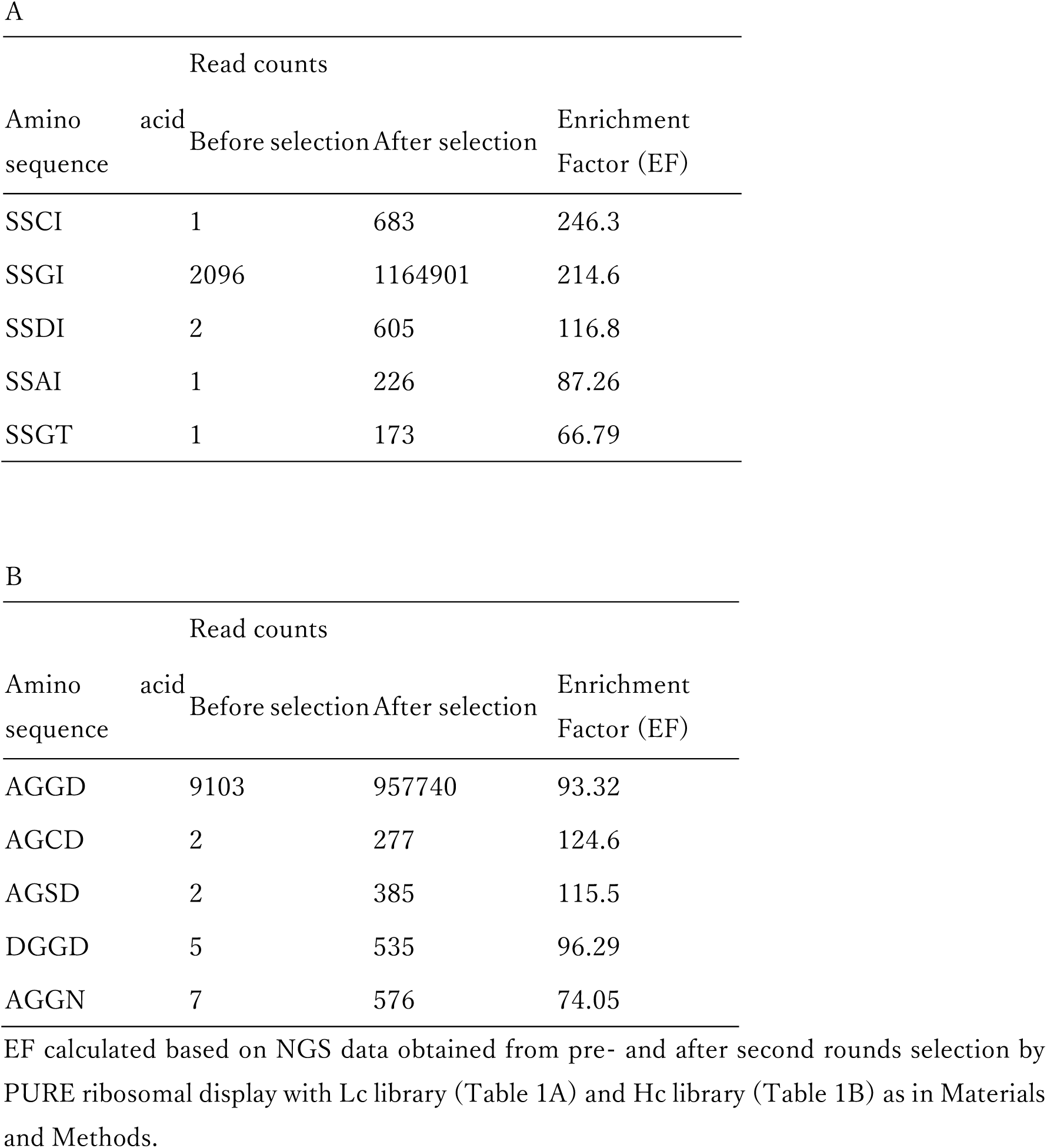
Top 5 EF in random region from NGS Analysis.

| Amino acid<br>sequence | Read counts |  | Enrichment<br>Factor (EF) |
| --- | --- | --- | --- |
|  | Before selection | After selection |  |
| SSCI | 1 | 683 | 246.3 |
| SSGI | 2096 | 1164901 | 214.6 |
| SSDI | 2 | 605 | 116.8 |
| SSAI | 1 | 226 | 87.26 |
| SSGT | 1 | 173 | 66.79 |

B
| Amino acid<br>sequence | Read counts |  | Enrichment<br>Factor (EF) |
| --- | --- | --- | --- |
|  | Before selection | After selection |  |
| AGGD | 9103 | 957740 | 93.32 |
| AGCD | 2 | 277 | 124.6 |
| AGSD | 2 | 385 | 115.5 |
| DGGD | 5 | 535 | 96.29 |
| AGGN | 7 | 576 | 74.05 |

### 3. Energy Calculation of anti-PA tag scFab variants

Results of energy minimization show that the Gibbs Free Energy of the A72D variant was the second lowest, next to the wild type, among the scFabs (wild type, A72D, G74C, G74S, and D75N) (Table 2). The G74S scFab had the highest Gibbs free energy value.

**Table 2.** Energy Minimization Calculation of complexes of PA tag and scFabs.

| Antibody Types | Gibbs Free Energy<br>(kJ/mol) |
| --- | --- |
| WT | $-7.27 \times 10^6$ |
| A72D | $-6.95 \times 10^6$ |
| G74C | $-6.93 \times 10^6$ |
| G74S | $-6.79 \times 10^6$ |
| D75N | $-6.86 \times 10^6$ |

Calculation was carried out as Materials and Methods.

**Table 3.** Binding ability of three anti-PA tag Fabs to PA tag measured by BLI assay.

| clone | $k_{on}$ (Ms<sup>-1</sup>) | $k_{on}$ Error | $k_{off}$ (s<sup>-1</sup>) | $k_{off}$ Error | $K_D$ (M) |
| --- | --- | --- | --- | --- | --- |
| WT | $2.686 \times 10^5$ | $4.721 \times 10^3$ | $5.532 \times 10^{-4}$ | $1.868 \times 10^{-5}$ | $2.06 \times 10^{-9}$ |
| G74C | $8.643 \times 10^4$ | $2.568 \times 10^3$ | $1.076 \times 10^{-3}$ | $3.914 \times 10^{-5}$ | $1.245 \times 10^{-8}$ |
| D75N | $1.552 \times 10^5$ | $1.066 \times 10^3$ | $2.499 \times 10^{-4}$ | $7.916 \times 10^{-6}$ | $1.611 \times 10^{-9}$ |

Three Fabs (WT, G74C, D75N) were expressed in *Brevibacillus choshinensis*, purified by IMAC and SEC, and then submitted to BLI assay.

**Table 4.**
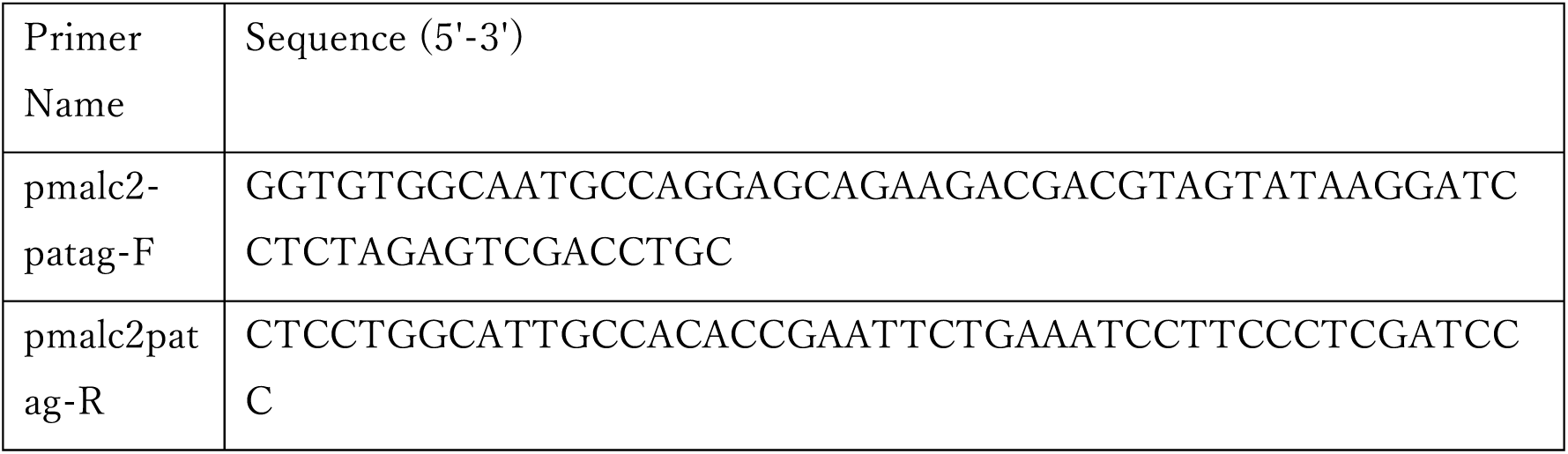
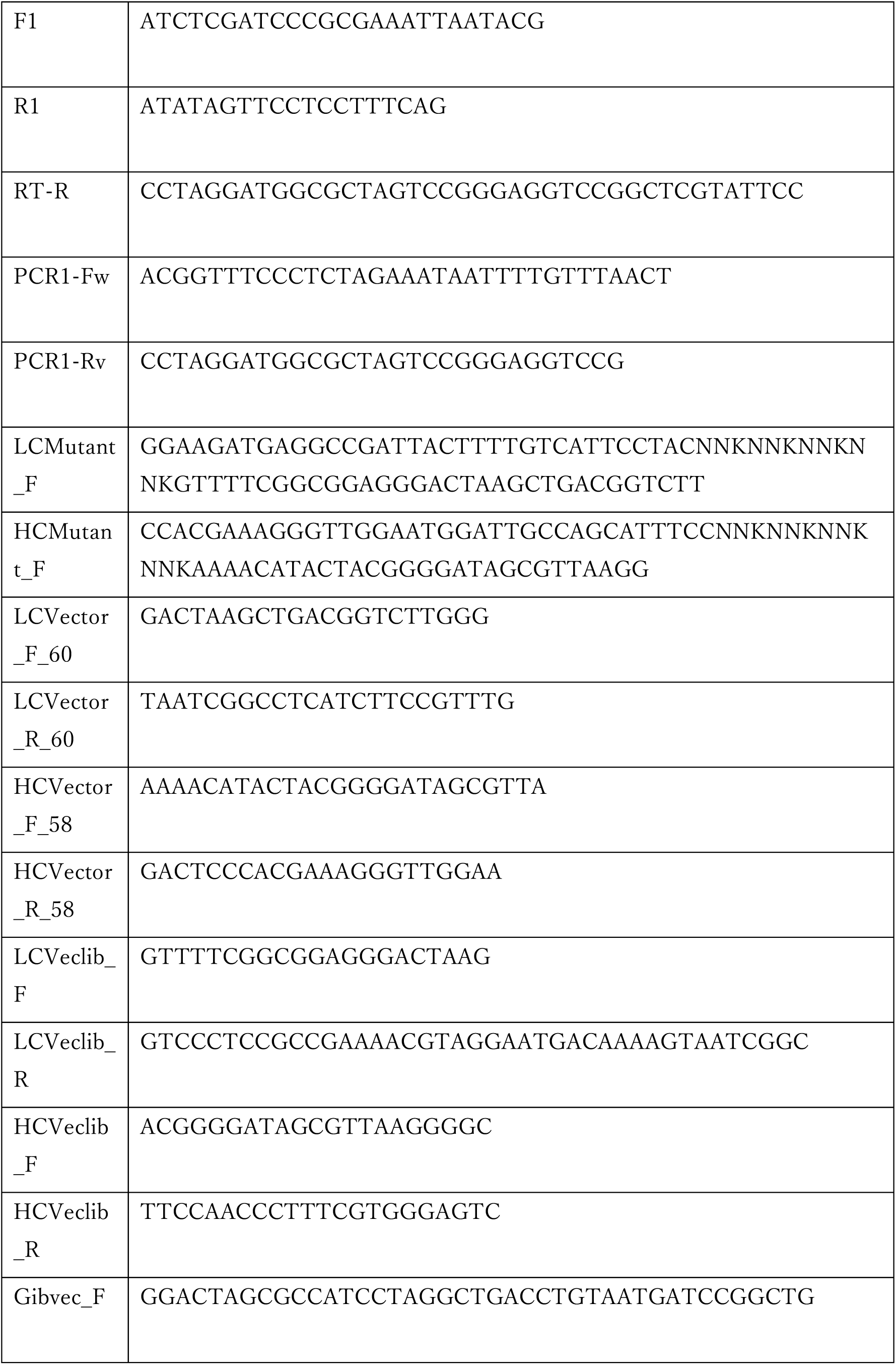
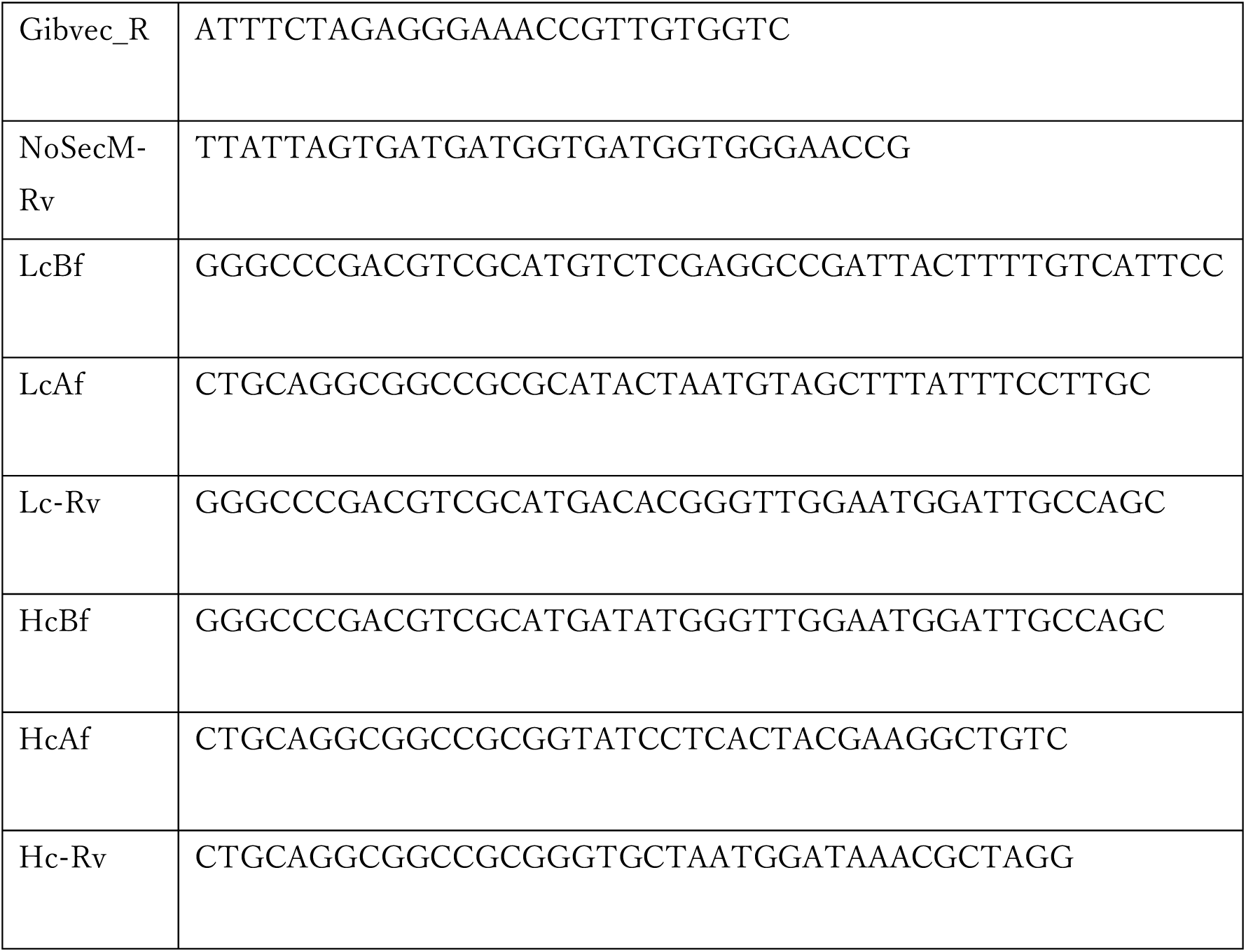
List of oligonucleotide primers.

### 4. Production of Fab variants Using *Brevibacillus choshinensis*

To evaluate the binding affinities of the anti-PA tag Fab variants obtained from the selection by PURE ribosome display, we produced Fab candidates using *Brevibacillus choshinensis*. Expression vectors for Fab were constructed, containing two genes encoding the light chain (VL-CL) and heavy chain (VH-CH) of Fab, each with its respective upstream secretion signal (Figure 6). The Fabs were purified from the culture supernatant by immobilized metal affinity chromatography (IMAC). The yields of wild-type, A72D, G74C, G74S, and D75N Fab variants were 1.24 mg, 1.40 mg, 1.53 mg, 1.78 mg, and 0.52 mg per 48 ml of culture supernatant respectively.

The binding activity of the partially purified Fabs were analyzed by ELISA, showing that among the four Fab variants (A72D, G74C, G74S, and D75N), G74C, G74S, D75N Fab variants had binding activity equal to or higher than the wild type. On the other hand, A72D variant had the lowest binding activity (Figure 5). The Gibbs Free Energy of G74S Fab variant is higher than G74C and D75N Fab variants. Then, G74C and D75N Fab variants were further purified using size exclusion chromatography (SEC).

**Figure 5.**
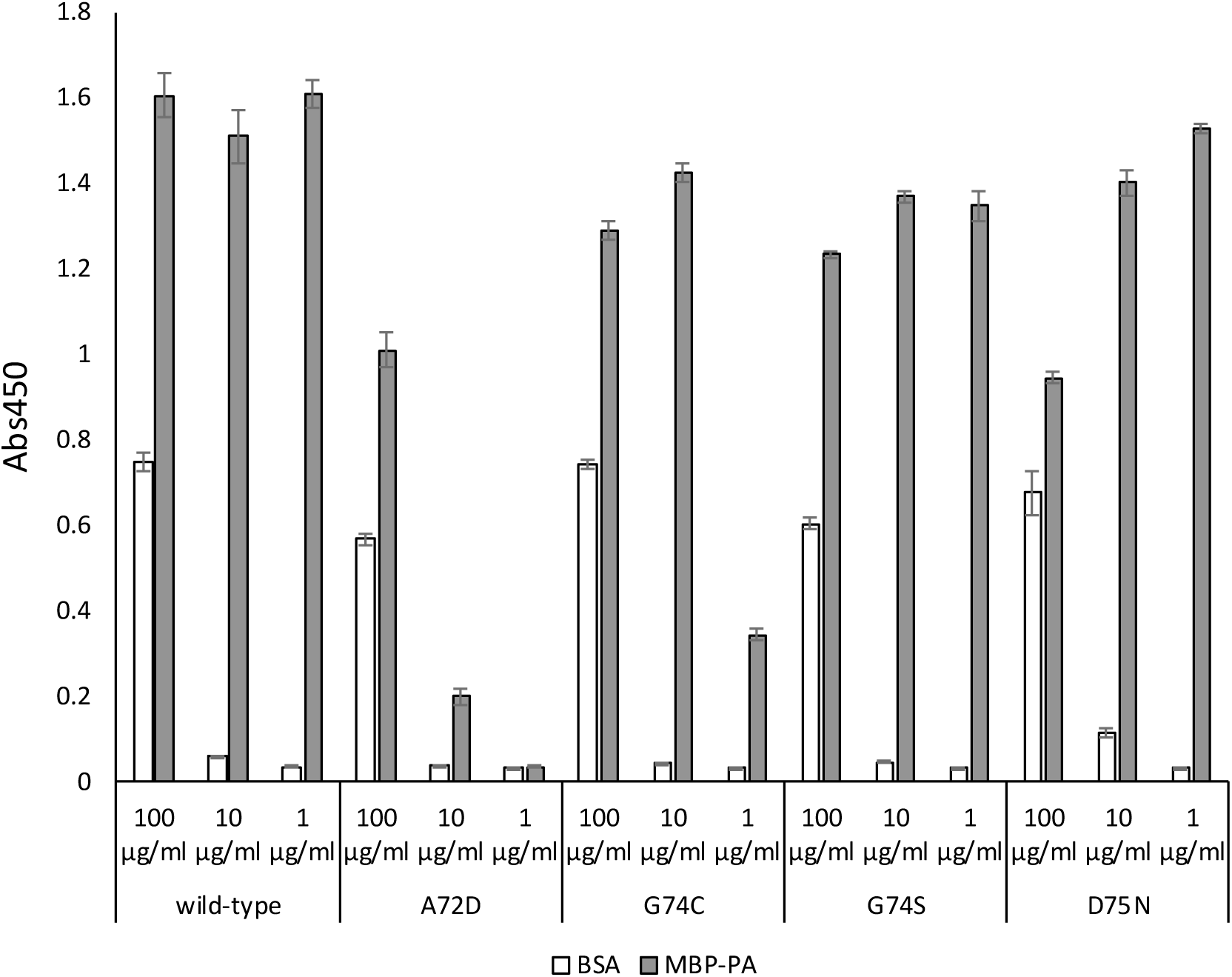
Evaluation of binding activity of Fabs expressed by *B. choshinensis* using ELISA. Fabs expressed by *B. choshinensis* were purified using immobilized metal affinity chromatography (IMAC) and diluted to concentrations of 100 µg/ml, 10 µg/ml, and 1 µg/ml. The diluted Fabs were then incubated in wells pre-coated with either 1% BSA or MBP-PA. X-axis: ELISA signal (Abs 450); error bars: standard deviation of the mean (n=3).

**Figure 6.**
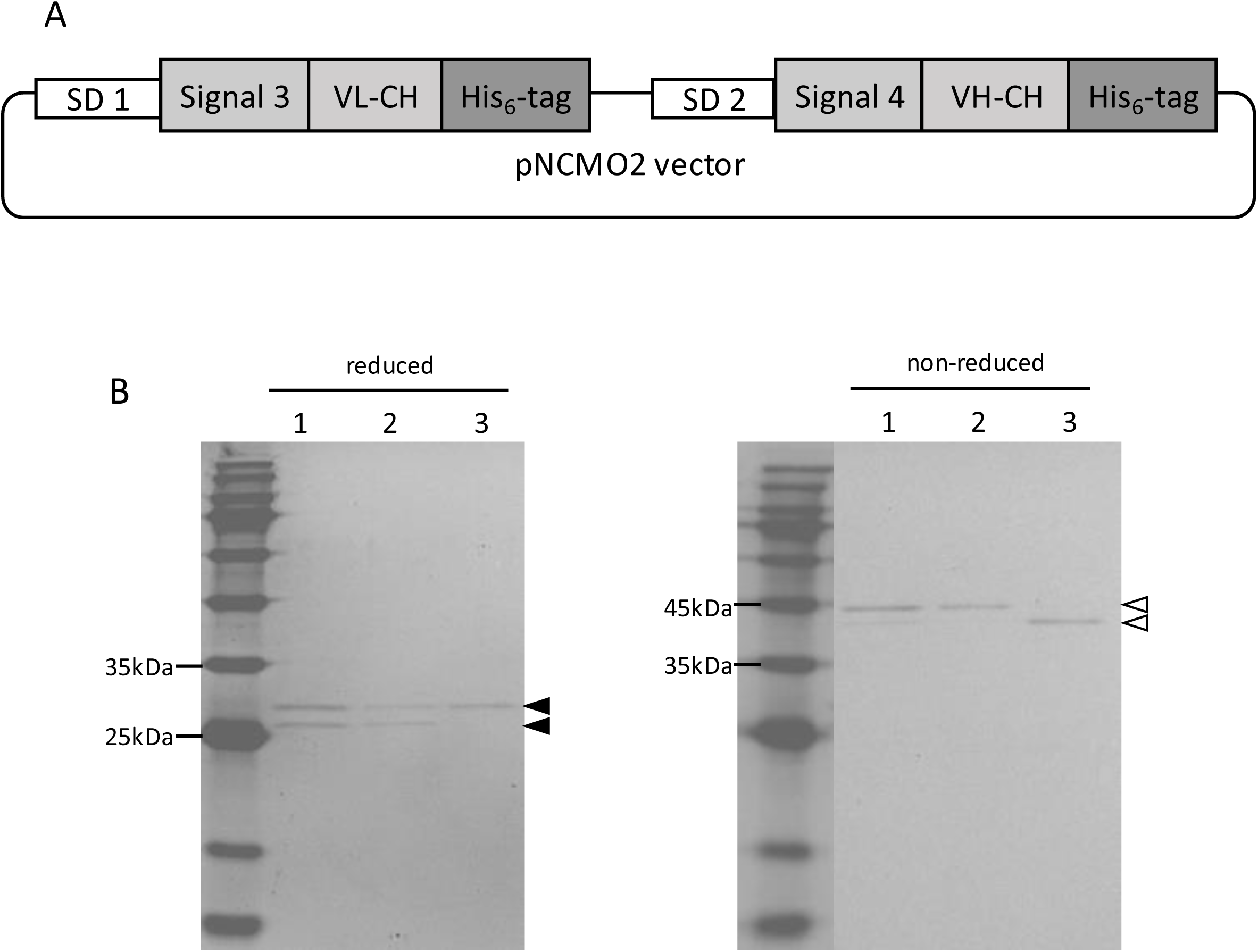
Expression of Fabs by *B. choshinensis* A: Illustration of expression vector of Fab for *B. choshinensis*. B: SDS–PAGE analysis of peak fractions with 12.5% SDS-PAGE gel. The peak fractions from SEC with a HiPrep 16/60 Sephacryl S-200 HR (Cytiva, Japan) were subjected to electrophoresis. Lane 1: WT Fab; Lane 2: D75N Fab variant; Lane 3: G74C Fab variant. Each samples were applied at 7 μl/lane for SDS-PAGE gel with and without a reducing agent. Black triangles indicate the Lc and Hc. White triangles indicate the heterodimer formed by Lc and Hc.

SDS-PAGE analysis of Fab variants purified by SEC under reducing conditions showed two bands with molecular weights consistent with Lc and Hc. Under non-reducing conditions, single band corresponding to heterodimers of Lc and Hc were observed. These results confirmed that wild-type anti-PA tag Fab, along with G74C and D75N Fab variants, were successfully purified and the disulfide bond between Lc and Hc was correctly formed. The final yield of Fab was approximately 0.09 mg per 48 ml of culture supernatant.

### 5. Analysis of Three Anti-PA Tag Fab variants Using BLI Assay

We investigated the molecular interactions between the purified Fab and MBP-PA using biolayer interferometry (BLI). The *K_D_* values for wild-type, G74C, and D75N Fab variants were 2.06 nM, 12.5 nM, and 1.61 nM, respectively (Figure 7).

**Figure 7.**
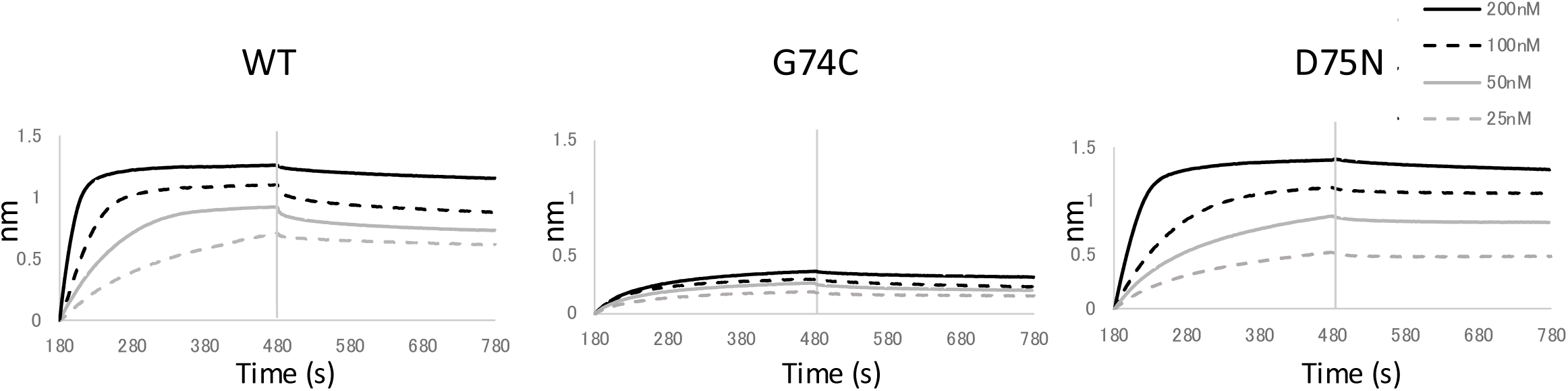
Kinetic analysis of anti-PA tag Fab using BLI assay APS sensors were sequentially dipped into MBP-PA (50 nM), each Fab (Gray dotted line: 25 nM, Gray solid line: 50 nM, Black dotted line: 100 nM, Black solid line: 200 nM) and PBS. Results obtained at 0 nM for each analyte were used as references. Data analysis and fitting were performed using BLItz Pro.

## Discussion

In this study, we developed a platform for rapidly and cost-effectively generating functional antibody variants by combining PURE ribosome display with NGS. The feasibility of this new PURE ribosome display method was demonstrated through the selection of both a model library with anti-PAscFab(118-120Ala) and a region-specific random library.

Our system is based on Fujino’s method with some modifications^10^. In this study, the SecM arrest sequence is used to form stable protein-ribosome-mRNA complexes. Additionally, we incorporated the upstream two amino acids, SK, from the SKIK sequence^26^, which is known to promote protein production, into the PURE ribosome display template^11,26,27^. Furthermore, we integrated a secretion expression system using *Brevibacillus* for detailed analysis of variants with enriched sequences.

The PURE ribosome display used here can be executed within seven hours per round. This includes two hours for In vitro transcription, 30 minutes for In vitro translation, one hour for PURE ribosome display, and two hours for reverse transcription and PCR. This is significantly faster compared to traditional methods like phage display. In the model library containing anti-PAscFab and anti-PAscFab(118-120Ala), the proportion of anti-PAscFab DNA increased greatly after one round of PURE ribosome display selection, indicating that the mRNA-ribosome-scFab ternary complex with high affinity for the PA tag was successfully enriched using PURE ribosome display.

The binding affinity of the antibody gene pool enriched by PURE ribosome display using the region-specific random library was evaluated by ELISA against PA-tagged MBP. As shown in Figure 4, the ELISA signal improved after selection in both the Lc and the Hc libraries. This indicated that the selection enriched genes for anti-PA tag Fab with high binding ability.

In the Hc library, NGS data analysis after two rounds of selection revealed that AGCD, AGSD, and DGGD had EF values of 124.63, 115.48, and 96.29, respectively, higher than the EF value of 93.32 for the wild-type sequence AGGD. The EF value for AGGN was 74.05. These results suggested that A72D, G74C, G74S, and D75N scFab variants might have superior binding abilities compared to the wild-type sequence.

Energy minimization by GROMACS showed that none of the variants were below the wild-type value. However, among the four variants, A72D had the lowest value and D75N had the second lowest value. Wild-type Fab, along with A72D, G74C, G74S, and D75N Fab variants were highly expressed using the *Brevibacillus choshinensis* secretion system. The Fabs were secreted into the culture medium and easily collected for purification. In accordance with previously published papers^28–30^, proline, Arg-HCl, and MgSO_4_ were added as folding aids.

The binding ability to the PA tag was evaluated by ELISA, and among G74C, G74S, and D75N Fab variants, which showed comparable activity to the wild type, the highest EF value and binding activity were observed for the D75N Fab variants. Further analysis by SDS-PAGE revealed an imbalance in the ratio of Hc to Lc. Therefore, wild-type, G74C, and D75N Fabs were further purified by SEC to resolve the imbalance between Hc and Lc (Figure 6). Approximately 0.09 mg of purified Fab per 48 ml of TM medium was obtained. This yield was lower than the previously reported yield of 1 mg per 96 ml, although direct comparison is not possible due to differences in the genes used.

The binding affinity of purified wild-type, G74C, and D75N Fabs to the PA tag was assessed by BLI assay. The binding affinity (*K_D_* = 12 nM) of the G74C Fab variant was over 5.7 times lower than that of the wild-type anti-PA tag Fab. The binding affinity (*K_D_* = 1.6 nM) of the D75N Fab variant was 1.3 times higher than that of the wild-type anti-PA tag Fab (*K_D_* = 2.1 nM). The *k_off_* (s^−1^) value for the D75N Fab variant was the smallest at 2.5×10-^4^ s^−1^, showing a 2.2 times difference compared to the wild-type Fab (5.5×10^−4^ s^−1^). This suggests that the D75N Fab gained the characteristic of being more resistant to dissociation from the PA tag after binding.

In conclusion, using PURE ribosome display, antibodies that bind to specific peptides were efficiently selected within seven hours per round. Combining NGS analysis allowed for the rapid and accurate collection of large amounts of sequence data, eliminating the need for laborious cloning steps. Furthermore, using the *Brevibacillus choshinensis* secretion system enabled simple collection and analysis of large quantity of Fab.

## Acknowledgments

Monami Kihara is supported by THERS Make New Standards Program for Next Generation Researchers and the CIBoG WISE Program. This research was financially supported in part by Grants-in-Aid for Scientific Research (No. 22520782 19105921) of the Japan Society for the Promotion of Science (JSPS), and by the Institute for Fermentation, Osaka (IFO): IFO research grant G-2022-3-021. The authors would like to thank Gene Frontier Corporation, Japan, for providing the PUREfrex components. We also extend our gratitude to Prof. Kohei Tsumoto and Dr. Ryo Matsunaga for their support in protein expression using Brevibacillus choshinensis.

## Notes

### Competing Interest Statement

The authors have declared no competing interest.

